# Loss of FMRP leads to translationally relevant functional connectivity differences in a rat model of Fragile X Syndrome

**DOI:** 10.64898/2026.08.28.747531

**Authors:** Jingjing Ye, Joanna A.B. Surl, Andrew G. McKechanie, Owen Dando, Milou Straathof, Rick M. Dijkhuizen, Andrew C. Stanfield, Sally M. Till, Peter C. Kind

**Affiliations:** Patrick Wild Centre, The University of Edinburgh; Edinburgh, UK; Simons Initiative for the Developing Brain; Edinburgh, UK; Institute for Neuroscience and Cardiovascular Research, The University of Edinburgh; Edinburgh, UK; Centre for Clinical Brain Sciences, The University of Edinburgh; Edinburgh, UK; UK Dementia Research Institute at The University of Edinburgh; Edinburgh, UK; Translational Neuroimaging Group, Center for Image Sciences, University Medical Center Utrecht and Utrecht University; Utrecht, The Netherlands

## Abstract

Fragile X syndrome (FXS), a leading monogenic cause of intellectual disability and autism-related features, results from loss of fragile X messenger ribonucleoprotein (FMRP). Although early synaptic and cellular abnormalities associated with the loss of FMRP are well described, it remains unclear how these changes shape the maturation of large-scale functional networks, or whether early pharmacological intervention can normalize circuit development. Our previous work demonstrated that cognitive deficits in *Fmr1^-/y^* rats emerge during development and can be prevented by brief early-life lovastatin treatment. Here, we asked whether large-scale functional connectivity (FC) shows a similarly dynamic developmental trajectory and whether early intervention alters its emergence. Using longitudinal resting-state functional magnetic resonance imaging (rsfMRI), we found that *Fmr1^-/y^* rats displayed an age-dependent FC phenotype, with increased connectivity within the retrosplenial cortex (RSC) at 4 weeks but reduced connectivity within the RSC and distributed brain networks by adulthood compared with wild-type controls. This suggests FC abnormalities emerge over development rather than representing a stable deficit. In contrast to its effects on cognitive measures, brief early-life lovastatin treatment did not prevent the emergence of connectivity abnormalities. Reduced RSC FC was also observed in a small cohort of individuals with FXS (n = 5 per group), supporting further investigation of functional connectivity measures alongside behavioural and molecular endpoints in translational studies of FXS.

**ONE SENTENCE SUMMARY:** An age-dependent brain connectivity phenotype in a rat model of FXS is not rescued by lovastatin and aligns with human RSC hypoconnectivity

## INTRODUCTION

Fragile X syndrome (FXS) is a leading monogenic form of intellectual disability and common cause of autism-related features. It is characterized by impairments in attention and executive function, sensory hypersensitivity, social anxiety and atypical social-communication (*1*), implicating disruption of distributed brain systems supporting higher-order cognition. FXS is caused by epigenetic silencing of the *FMR1* gene, resulting in loss of the fragile X messenger ribonucleoprotein, FMRP (*2, 3*). As an RNA-binding protein that regulates synaptic protein synthesis, FMRP is key for cell function at multiple levels including circuit maturation (*4*). In rodent models, loss of FMRP leads to widespread alterations in neuronal excitability, synaptic plasticity, and dendritic spine development across multiple brain regions accompanied by behavioural phenotypes relevant to FXS (*5–7*). These findings suggest that early synaptic dysfunction disrupts the development of large-scale brain networks. Consistent with this, human neuroimaging studies report altered functional connectivity (FC) in FXS, including abnormalities within canonical resting state networks such as the default mode network (DMN) (*8–10*).

We previously demonstrated reduced associative learning in two distinct rat models of FXS (*11, 12*). Importantly, the emergence of this phenotype can be prevented by brief, early life treatment with lovastatin (*12*), an HMG-CoA reductase inhibitor widely used to treat hypercholesterolemia (*13*). Improvements in associative learning as well as normalization of protein synthesis persisted after treatment withdrawal, suggesting that loss of FMRP alters developmental trajectories that may have long-lasting benefits if corrected early (*12*). Consistent with this, lovastatin has been shown to correct alterations in synaptic plasticity, excitability and seizure susceptibility in a mouse model of FXS (*14, 15*). However, whether such early interventions also normalize large-scale brain networks remains unclear; this question is particularly relevant given the mixed outcomes of lovastatin clinical trials, which have not established a clear therapeutic benefit (*16–18*). This raises the possibility that preclinical rescue of cellular and behavioural phenotypes may not extend to systems-level brain organisation relevant to FXS.

To address this question, we combined longitudinal and cross-species approaches to test whether FXS-related circuit abnormalities emerge and evolve across development and whether they are sensitive to early pharmacological intervention. We first characterized resting-state FC in adults and across development in a rat model of FXS to define network level abnormalities. We then tested whether early-life lovastatin treatment modifies large-scale network organization in *Fmr1* knockout (*Fmr1^-/y^)* rats. Finally, we examined whether these alterations are conserved in humans with FXS. Since resting state networks are conserved across species, including humans, non-human primates, rats and mice (*19*), this approach allows us to determine whether Fragile X-related network level abnormalities emerge over development, are conserved across species, and are sensitive to early pharmacological intervention.

## RESULTS

### RSC and global hypoconnectivity in adult *Fmr1^-/y^* rats

As brain-wide FC has not yet been characterized in *Fmr1^-/y^*rats, we used rsfMRI to map whole-brain networks in a cohort of lightly anaesthetized *Fmr1^-/y^* rats and their wild-type (WT) littermates. To identify network-level alterations, we first mapped whole-brain FC in 15-week-old rats using voxel-wise degree centrality (DC, (*20*)). This analysis revealed reduced DC in the retrosplenial cortex (RSC), a key node of the posterior DMN, in *Fmr1^-/y^*rats compared to WT littermates (Fig. 1A; p < 0.01, threshold-free cluster enhancement (TFCE)-corrected). To further characterize RSC connectivity, seed-based analyses were performed using the RSC subregion (Fig. 1B). *Fmr1^-/y^* rats showed reduced connectivity relative to WT within the DMN and more broadly distributed networks (Fig. 1C-F). Consistent with these maps, quantification of FC strength demonstrated significantly reduced local connectivity within the DMN (Fig. 1G; p < 0.05) and reduced global connectivity across the brain in *Fmr1^-/y^* rats (Fig. 1H; p < 0.05).

**Fig. 1.**
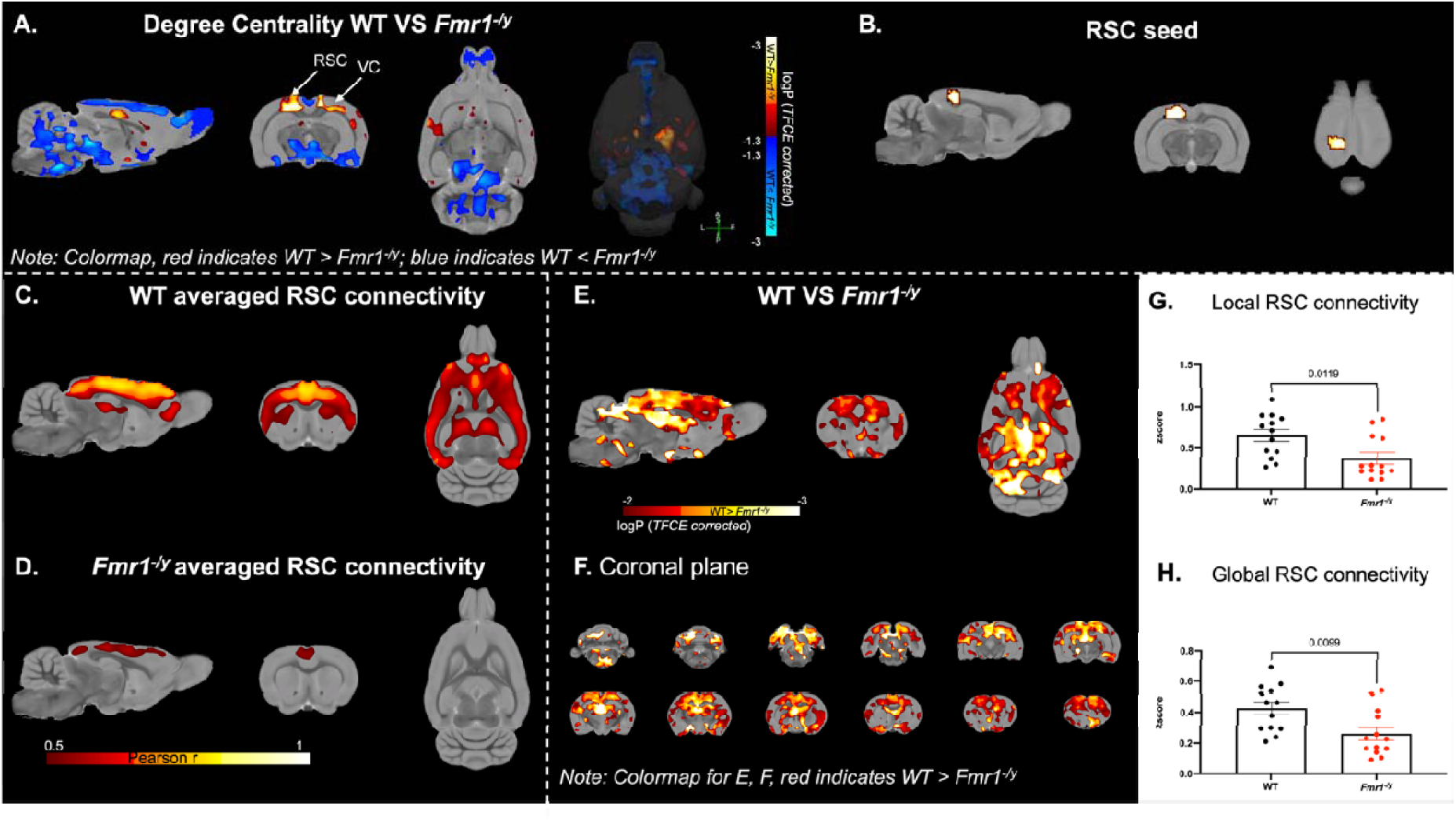
Reduced local and global RSC connectivity in *Fmr1^-/y^* rats. (**A**) Degree centrality (DC) analysis at 15 weeks of age, showing reduced RSC DC (red voxels) and increased cerebellar DC (blue voxels) in *Fmr1^-/y^* rats compared to WT controls. (**B**) RSC seed defined from the DC analysis. (**C, D**) Group averaged RSC functional connectivity maps for *Fmr1^-/y^* and WT rats displayed at r > 0.5. (**E, F**) Voxel-wise comparison of RSC connectivity, with regions showing greater connectivity in WT than *Fmr1^-/y^*shown in red. (**G, H**) Quantification of local and global RSC functional connectivity, expressed as z-transformed Pearson correlation coefficients. Data shown as mean ± SEM. Group differences were assessed using two-sample t-test. Group size: WT, n = 13; *Fmr1^-/y^*, n = 13. For visualisation, group averaged connectivity maps were thresholded at r > 0.5 to emphasise robust cortical connectivity patterns. Voxel-wise group comparisons were performed using TFCE-correction (p<0.01). Abbreviations: RSC, retrosplenial cortex; VC, visual cortex; SEM, standard error mean; WT, wild-type; TFCE, threshold-free cluster enhancement.

To determine whether these RSC abnormalities reflected broader network disruption, we next assessed resting-state FC across 25 atlas-defined regions of interest (ROIs) spanning cortical, limbic, and cerebellar regions (Fig. 2A). Group-averaged connectivity matrices showed reduced global connectivity in *Fmr1^-/y^* rats relative to WT controls (Fig. 2B), with statistical comparison confirming prominent reductions in cortico-limbic connections (Fig. 2C; network-based statistic (NBS)-corrected, p < 0.05).

**Fig. 2.**
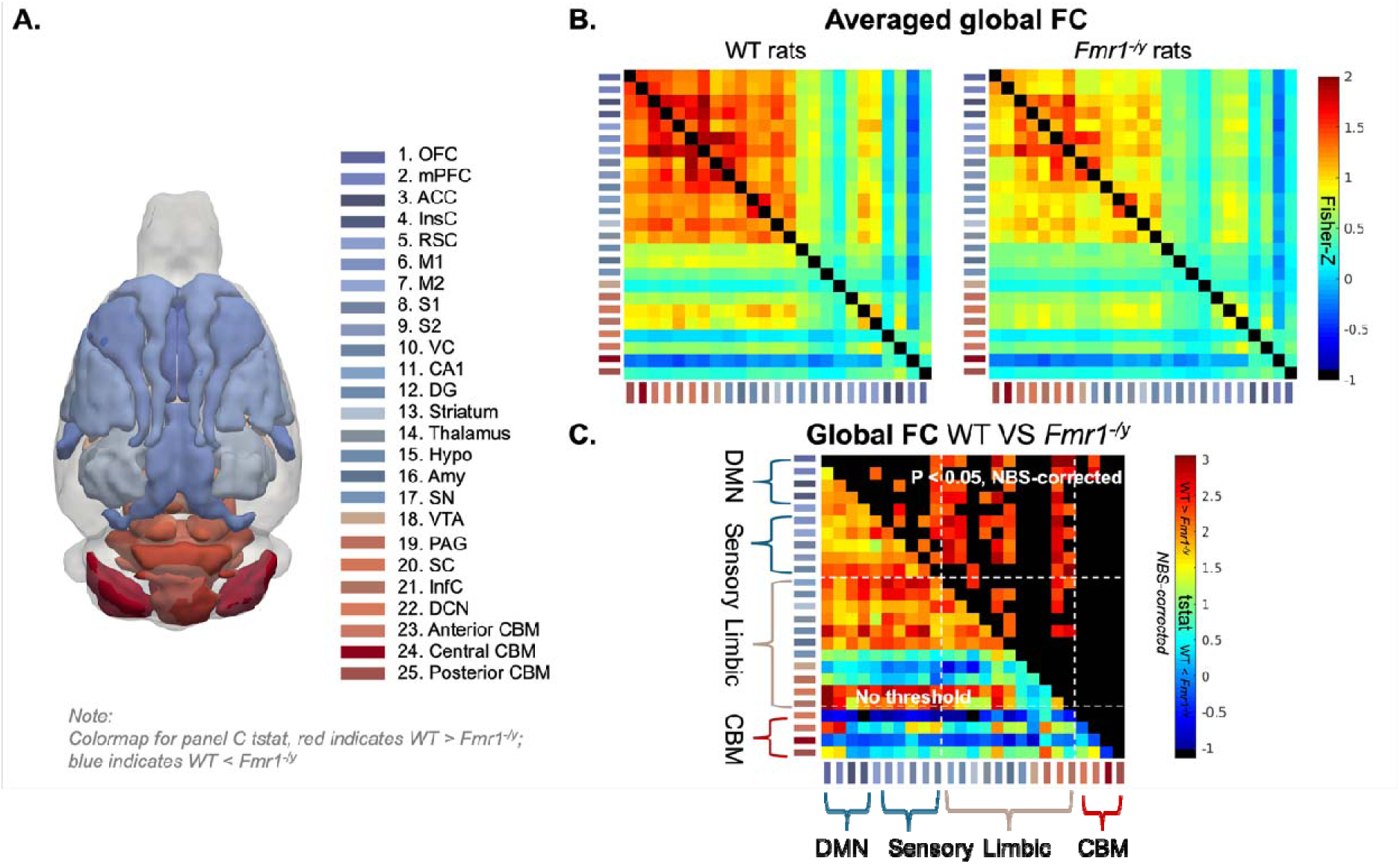
Reduced global functional connectivity in *Fmr1^-/^*^y^ rats. (**A**) Regions of interest (ROIs) and colour coding for the 25 anatomical brain regions included in the functional connectivity (FC) analysis. (**B**) Group-averaged global FC matrices for WT and *Fmr1^-/y^*rats. (**C**) t-statistic matrix showing between-group differences in FC (red, WT > *Fmr1^-/y^*; blue, WT < *Fmr1^-/y^*), identified using NBS with two-sample t-tests. White dashed lines indicate boundaries between cortical (DMN and sensory), subcortical (limbic), and cerebellar networks. Group sizes: WT, n = 13; *Fmr1^-/y^*, n = 13. Brain regions: 1. OFC, orbitofrontal cortex; 2. mPFC, medial prefrontal cortex; 3. ACC, anterior cingulate cortex; 4. InsC, insular cortex; 5. RSC, retrosplenial cortex; 6. M1, primary motor cortex; 7. M2, secondary motor cortex; 8. S1, primary somatosensory cortex; 9. S2, secondary somatosensory cortex; 10. VC, visual cortex; 11. CA1, hippocampal CA1 region; 12. DG, dentate gyrus; 13. Striatum; 14. Thalamus; 15. Hypo, hypothalamus; 16. Amy, amygdala; 17. SN, substantia nigra; 18. VTA, ventral tegmental area; 19. PAG, periaqueductal gray; 20. SC, superior colliculus; 21. InfC, inferior colliculus; 22. DCN, deep cerebellar nuclei; 23. Anterior CBM, anterior cerebellum; 24. Central CBM, central cerebellum; 25, Posterior CBM, posterior cerebellum.: Cerebellar subdivisions were defined according to {Apps, 2009 #1285; Ciapponi, 2023 #1286}: anterior cerebellum (lobules I-V), central cerebellum (lobules VI-VII), posterior cerebellum (lobules VIII-X, and Crus I/II). Abbreviations: FC, functional connectivity; DMN, default mode network; NBS, Network-Based Statistic.

### Altered developmental trajectory of functional connectivity in *Fmr1^-/y^* rats

Because the RSC is a core node of the DMN, and DMN connectivity has been reported to show age-dependent alterations in ASD (*21–26*), we tested whether RSC brain-wide connectivity shows an altered developmental trajectory in *Fmr1^-/y^* rats. To address this, we performed longitudinal seed-based connectivity analyses at 4, 8, 12, and 15 weeks of age, using the RSC subregion identified by our initial DC analysis as the seed (Fig. 1B). At 4 weeks, statistical comparison revealed significantly increased RSC connectivity in *Fmr1^-/y^*rats relative to WT controls, specifically with the insular cortex, medial prefrontal cortex, and caudate putamen (Fig. 3A; p < 0.01, TFCE-corrected). By 15 weeks, the early hyperconnectivity was reversed, with significant hypoconnectivity in *Fmr1^-/y^*rats (Fig. 3B and 1E). To track the full trajectory, averaged connectivity maps for WT and *Fmr1^-/y^* rats across all developmental timepoints, including 8 and 12 weeks, are shown as 3D glass brain projections (Fig. 3C). Longitudinal quantification of connectivity strength demonstrated an age-dependent shift from early hyperconnectivity to later hypoconnectivity in *Fmr1^-/y^* rats, with no significant group differences observed during the intermediate 8- and 12-week stages (Fig. 3D; p < 0.05).

**Fig. 3.**
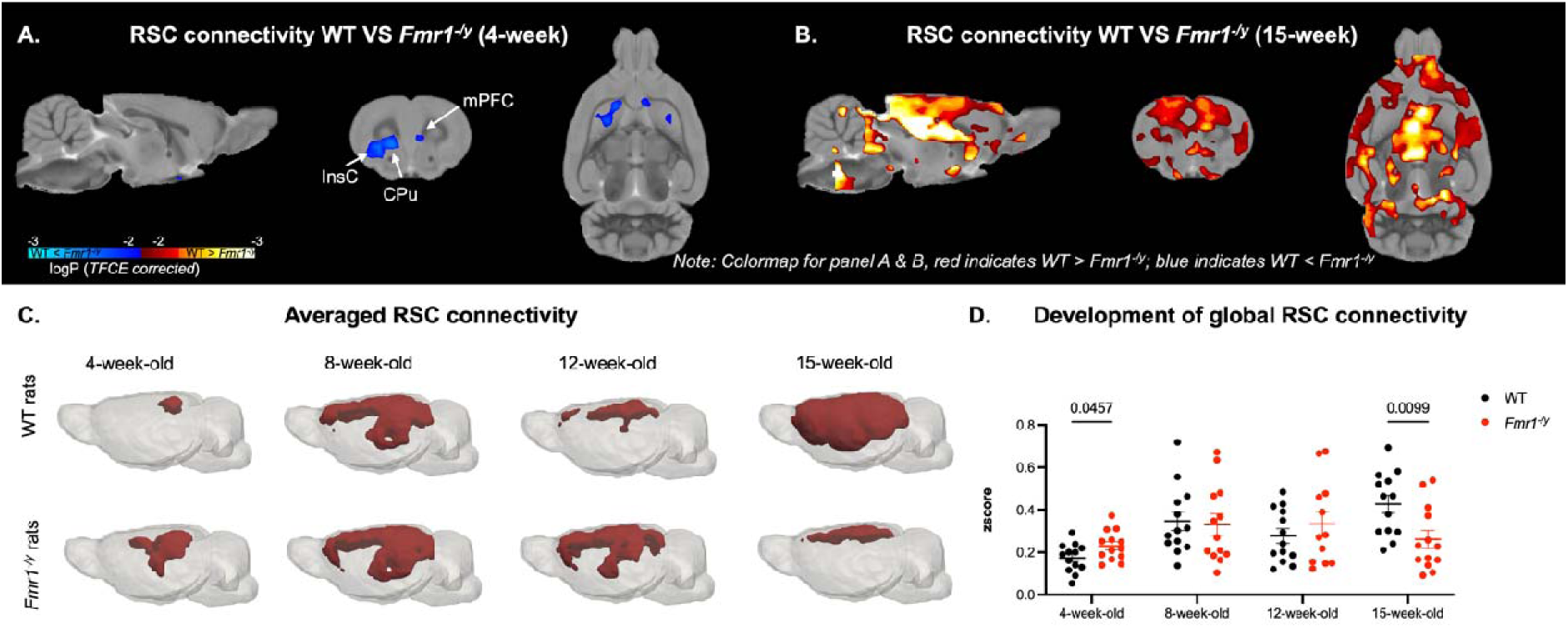
Developmental trajectory of RSC functional connectivity in WT and *Fmr1^-/y^* rats. (**A**) Voxel-wise comparison of RSC FC at 4 weeks of age. *Fmr1^-/y^*rats exhibited significantly greater connectivity than WT controls (blue), particularly with regions of the mPFC and InsC (TFCE-corrected, p < 0.01). (**B**) At 15 weeks, *Fmr1^-/y^* rats showed widespread reductions in RSC connectivity relative to WT controls (red), involving cortical, subcortical and hindbrain regions, including the PAG and cerebellum (TFCE-corrected, p < 0.01). (**C**) Group-averaged RSC FC maps at 4, 8, 12, and 15 weeks for WT and *Fmr1^-/y^* rats, displayed at r > 0.5. (**D**) Quantification of global RSC FC expressed as z-transformed Pearson correlation coefficients. Mixed-effects analysis revealed a significant main effect of age (F_(2.895, 68.52)_ = 6.964, p = 0.0004) and a significant age × genotype interaction (F_(3, 71)_ = 4.309, p = 0.0075), but no main effect of genotype (F_(1, 24)_ = 0.2419, p = 0.6273). Data are shown as mean ± SEM. Group sizes: WT, 4-15 weeks, n = 13; *Fmr1^-/y^*, 4, 8, and 15 weeks, n = 13 and 12 weeks, n = 12. Abbreviations: CPu, caudate putamen; mPFC, medial prefrontal cortex; InsC, insular cortex; DMN, default mode network; PAG, periaqueductal gray; RSC, retrosplenial cortex; SEM, standard error of the mean; TFCE, threshold-free cluster enhancement; WT, wild type.

We next examined whether the developmental shift observed in RSC connectivity was also evident at the level of whole-brain FC. FC matrices were computed for *Fmr1^-/y^* and WT rats at 4, 8, 12, and 15 weeks of age (Fig. 4). Linear mixed-effects modelling of global FC revealed a significant genotype × age interaction (p < 0.0001), as well as a main effect of age (p < 0.0001). Consistent with the seed-based approach, a post hoc simple effects analysis (Table 1) demonstrated a significant genotype difference at 15 weeks, with *Fmr1^-/y^* rats exhibiting reduced global FC compared to WT controls (T = 4.933, p = 0.0001). To validate this trajectory, we additionally modeled the first principal component (PC1) derived from a principal component analysis (PCA; explaining 77.78% of the total variance) further. Mixed-effects modelling of PC1 scores indicated a significant effect of age (Fig. S1 A,B), consistent with a robust developmental shift in global FC across timepoints.

**Fig. 4.**
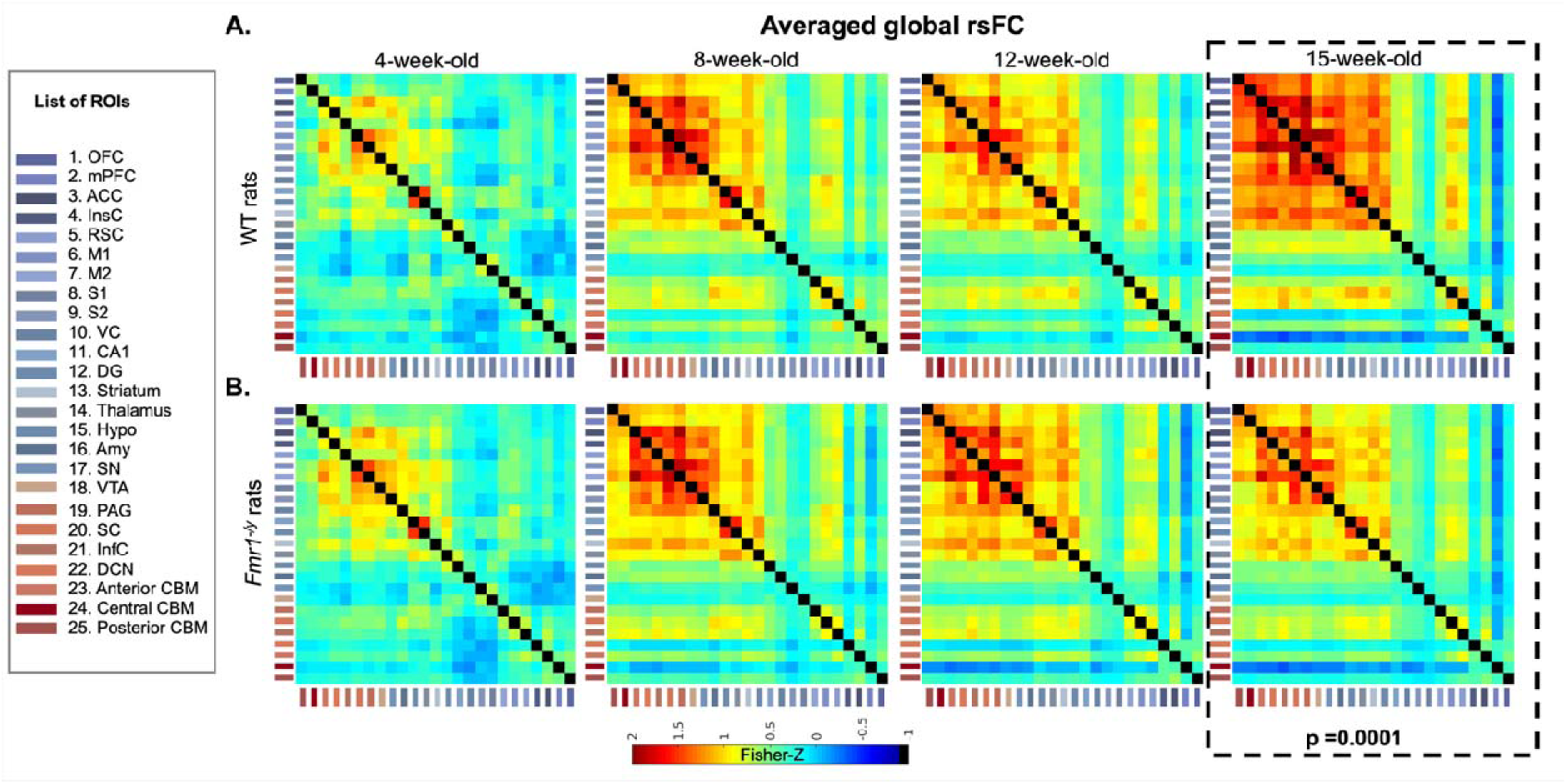
Developmental trajectories of global ROI-based FC in WT and *Fmr1^-/y^* rats. (**A**) Averaged FC matrices for WT and (**B**) *Fmr1^-/y^* rats at each developmental timepoint (4, 8, 12, 15 weeks). Group size: WT, 4-15 weeks, n = 13; *Fmr1^-/y^*, 4, 8, 15 weeks, n = 13; 12 weeks, n = 12. Mixed-effects analysis: main effect of age (F_(3,23094)_ = 1193.45, *p* < 0.0001); main effect of genotype (F_(1,24)_ = 1.76, *p* = 0.1976); age × genotype interaction (F_(3,23094)_ = 147.40, *p* < 0.0001). Abbreviations: FC, functional connectivity; SEM, standard error mean; WT, wild- type. Note: List of ROIs are provided on the right side of the figure, and abbreviations for brain regions are provided in the Figure 2 legend.

**Table 1.** Participants demographics.

| Characteristics | FXS group | Control groups |
| --- | --- | --- |
| Number | 5 (4 males 1 female) | 5 (male) |
| Age, mean (SD) | 18.6 (0.5) | 28 (3.08) |
| IQ, mean (SD) | 60.2 (12.5) | 114.5 (14.33) |
| ADOS-2 C+SI total score (SD) | 16 (4.2) | / |

### Early lovastatin treatment does not rescue hypoconnectivity in *Fmr1^-/y^* rats

We previously identified that the time window between 4 and 9 weeks of age is critical for the development of mPFC-dependent circuits underlying associative memory (*12*). Brief lovastatin administration during this period prevented the emergence of deficits in associative memory, as well as synaptic plasticity deficits in the mPFC of *Fmr1^-/y^* rats, with long-lasting effects on behaviour and protein synthesis (*12*). Given that both mPFC and RSC are key components of the DMN, we investigated whether brief, early lovastatin intervention between 4 weeks and 9 weeks of age would normalize FC across development in *Fmr1^-/y^*rats. To test this, *Fmr1^-/y^* rats were assigned to either lovastatin-enriched or control diet groups after the imaging scan at 4 weeks of age, and RSC-centered DMN connectivity as well as global FC were assessed at 8 and 15 weeks of age, corresponding to 4 weeks after treatment onset and 6 weeks after cessation.

RSC connectivity was analyzed using the same seed region defined in Fig. 1B, with connectivity patterns visualized as 3D glass brain projections (Fig. 5A,B) and connectivity strength quantified across groups (Fig. 5C). Mixed-effects modelling revealed a significant main effect of genotype (p < 0.05), but no significant effects of age, treatment, or interactions. These findings indicate that lovastatin administration during the developmental window previously associated with behavioural and synaptic rescue failed to rescue RSC hypoconnectivity phenotype in *Fmr1^-/y^* rats.

**Fig. 5.**
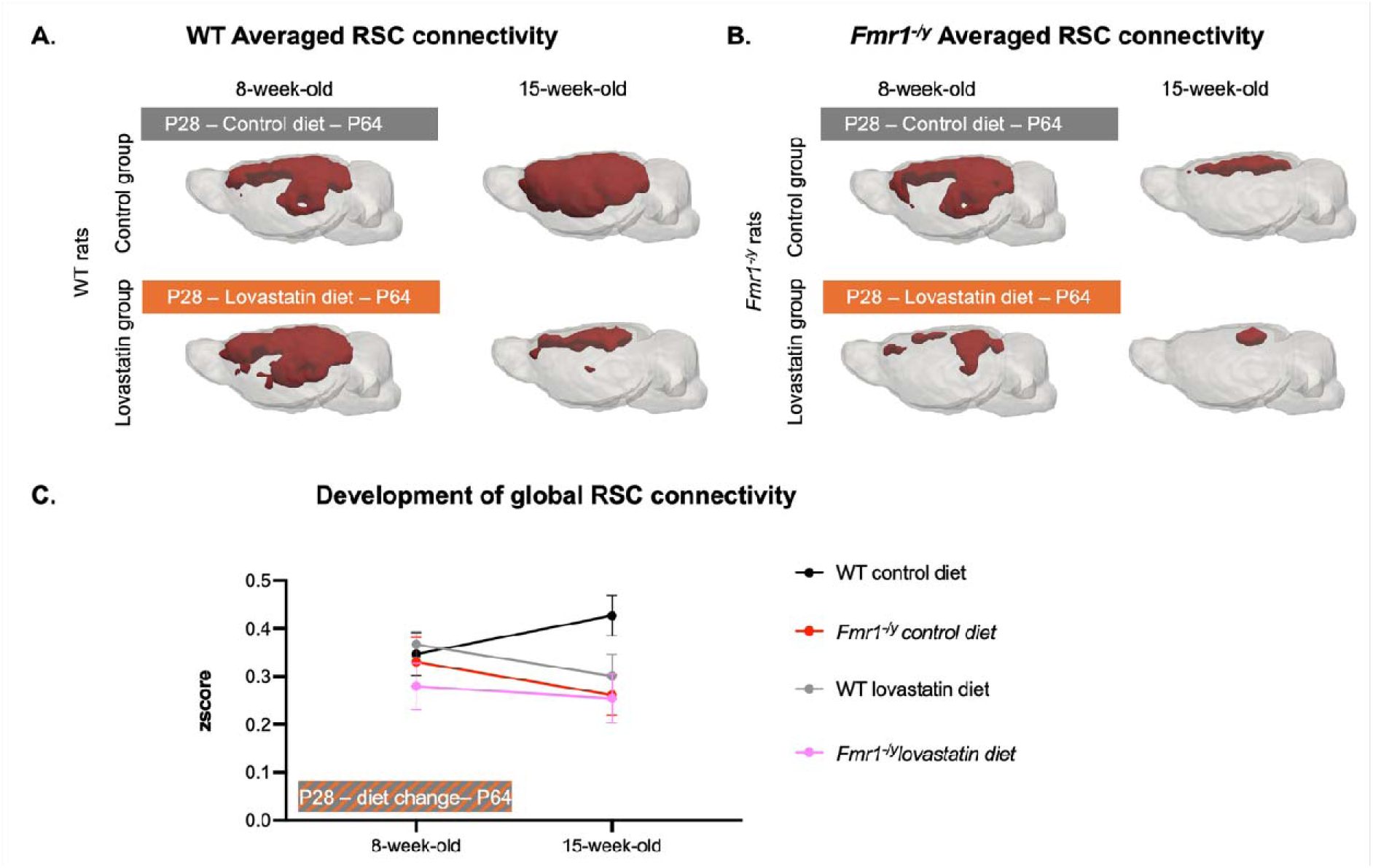
Development of RSC functional connectivity following early lovastatin treatment in WT and *Fmr1^-/y^*rats. **(A, B**) Group-averaged RSC seed-based FC maps for WT and *Fmr1^-/y^*rats under control and lovastatin diets at 8 and 15 weeks of age. Maps are displayed r > 0.5. (**C**) Longitudinal change in RSC FC from 8 to 15 weeks in WT and *Fmr1^-/y^* rats with or without lovastatin treatment intitiated after the 4-week scan. Data shown as mean ± SEM. Group sizes: WT control diet, n = 13; *Fmr1^-/y^* control diet, n = 13; WT Lovastatin diet, n = 14; *Fmr1^-/y^* Lovastatin diet, n = 11 (8 weeks) and n = 12 (15 weeks). Mixed-effects analysis revealed a main effect of genotype (F_(1,50)_ = 5.96, p = 0.02), but no main effects of age (F_(1,50)_ = 0.34, p = 0.56) or treatment (F_(1,45)_ = 2.00, p = 0.16) and no significant genotype × treatment, genotype × age, or age × genotype × treatment interactions (all p > 0.05). Abbreviations: RSC, retrosplenial cortex; SEM, standard error mean; WT, wild-type.

To assess whether RSC effects reflected broader network-level changes, we examined global FC correlation matrices across development and treatment conditions (Fig. 6). Linear mixed-effects analysis of global FC revealed significant genotype × age × treatment (p < 0.0001), age × treatment (p < 0.0001), as well as a main effect of age (p < 0.0001) and genotype (p < 0.05). Post hoc comparisons (Table 2) demonstrated a significant genotype effect at 15 weeks, with WT rats on the control diet exhibiting higher global FC than *Fmr1^-/y^* rats receiving either the control or lovastatin diet (WT_control diet_ > *Fmr1^-/y^*_control diet_, T= 3.227, p = 0.0135; WT_control diet_ > *Fmr1^-/y^*_lovastatin diet_, T = 2.671, p = 0.0291), suggesting that brief lovastatin treatment did not prevent the emergence of global FC deficits in *Fmr1^-/y^* rats. Interestingly, lovastatin-treated WT rats also exhibited reduced global FC compared with WT controls at 15 weeks of age (T = 2.535, p = 0.0291), indicating that lovastatin may negatively impact FC in otherwise typical brains. To determine whether a treatment effect could be captured in a lower-dimensional representation of the data, we also performed PCA on the global connectivity matrices; the resulting PC1 accounted for 77.20% of the total variance. Mixed-effects analysis of PC1 scores did not reveal any evidence that lovastatin altered the trajectory of global FC in *Fmr1^-/y^* rats (Fig. S1C).

**Fig. 6.**
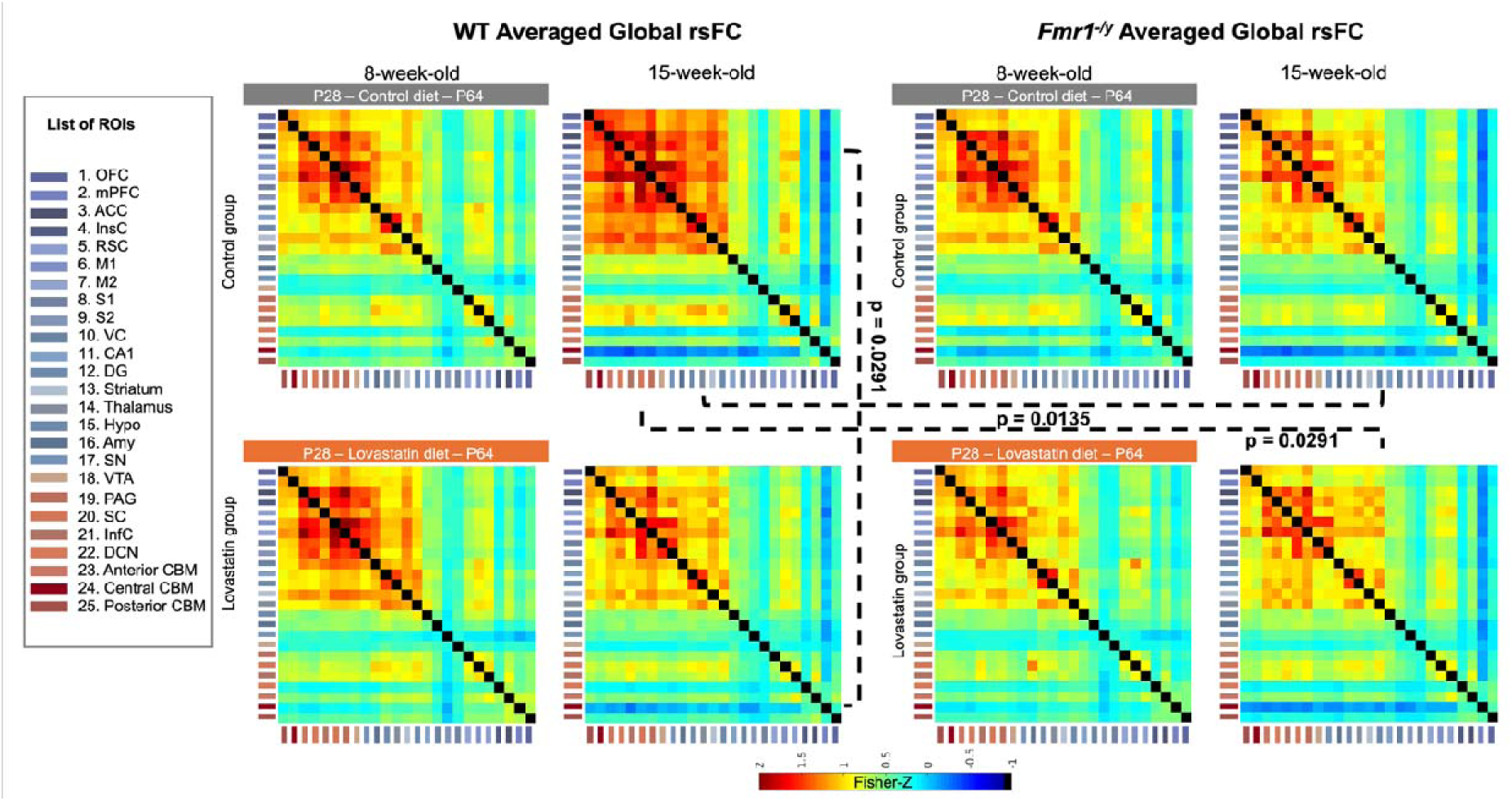
Changes in global rsFC following application of lovastatin diet from 5 to 9 weeks of age. Group-averaged global FC matrices of WT and *Fmr1^-/y^* rats under control (upper) and lovastatin diet (lower) diet conditions. Group sizes: WT control diet, n = 13 (8 and 15 weeks); *Fmr1^-/y^* control diet, n = 13 (8 and 15 weeks); WT lovastatin diet, n = 14 (8 and 15 weeks); *Fmr1^-/y^* lovastatin diet, n = 11 (8 weeks) and n = 12 (15 weeks). Linear mixed-effects analysis revealed a main effect of age (F_(1, 15296)_ = 59.92, p < 0.0001), a main effect of genotype (F_(1,48)_ = 5.15, p = 0.0278), and a significant age x treatment interaction (F_(1, 15296)_ = 59.48, p < 0.0001), as well as a significant genotype x age x treatment interaction (F_(1,15296)_ = 104.39, p < 0.0001). There was no main effect of treatment (F(_1,48_) = 0.51, p = 0.48), nor significant age × genotype (F(_1,15296_) = 2.35, p = 0.12) or genotype × treatment (F(_1,48_) = 0.82, p = 0.37) interactions. Abbreviations: FC, functional connectivity; SEM, standard error mean; WT, wild-type. ROI abbreviations are listed in Figure 2.

**Table 2.**
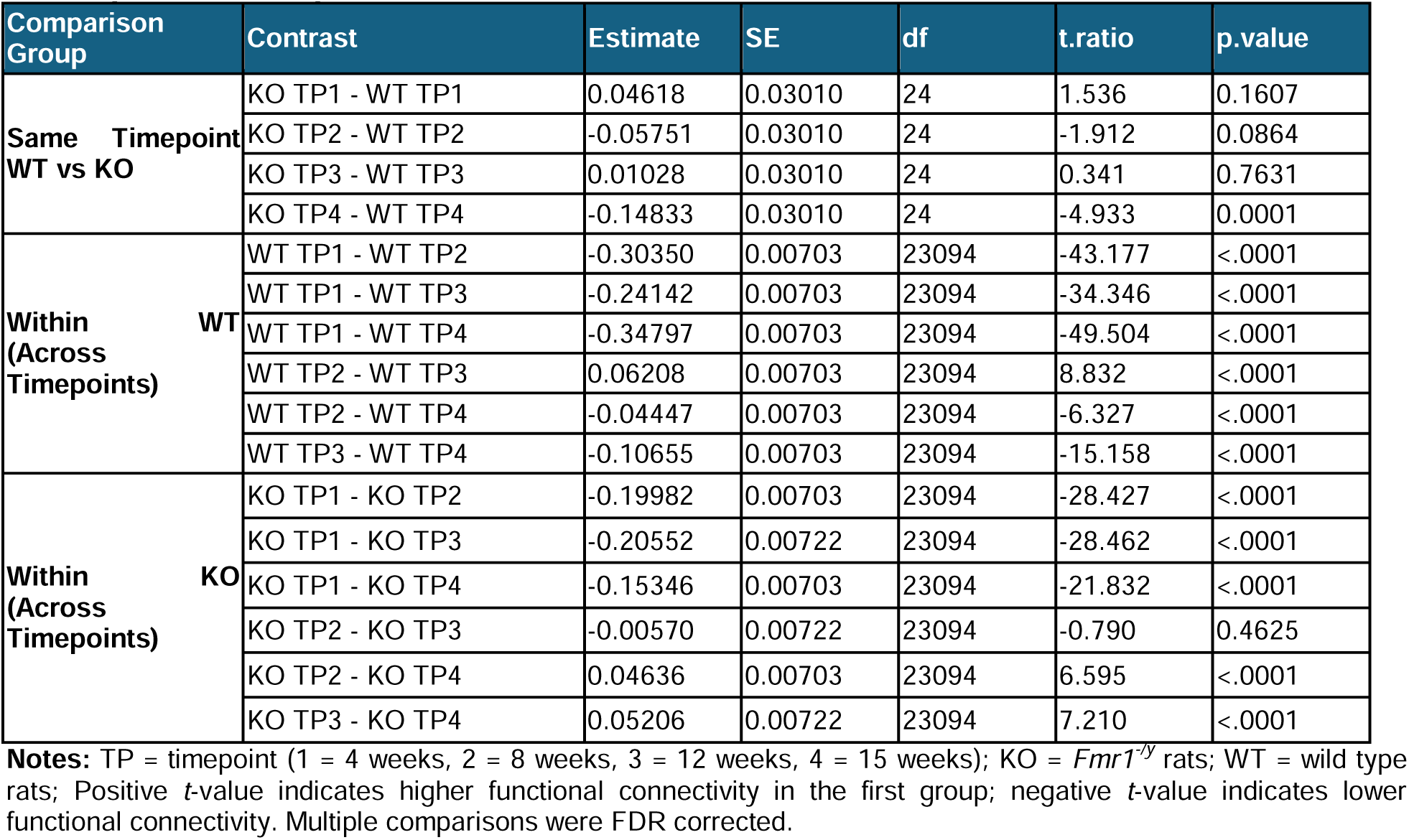
Post-hoc comparisons of global rsFC between genotypes and across developmental timepoints.

| Comparison Group | Contrast | Estimate | SE | df | t.ratio | p.value |
| --- | --- | --- | --- | --- | --- | --- |
| **Same Timepoint**<br>**WT vs KO** | KO TP1 - WT TP1 | 0.04618 | 0.03010 | 24 | 1.536 | 0.1607 |
|  | KO TP2 - WT TP2 | -0.05751 | 0.03010 | 24 | -1.912 | 0.0864 |
|  | KO TP3 - WT TP3 | 0.01028 | 0.03010 | 24 | 0.341 | 0.7631 |
|  | KO TP4 - WT TP4 | -0.14833 | 0.03010 | 24 | -4.933 | 0.0001 |
| **Within (Across Timepoints)**<br>**WT** | WT TP1 - WT TP2 | -0.30350 | 0.00703 | 23094 | -43.177 | <.0001 |
|  | WT TP1 - WT TP3 | -0.24142 | 0.00703 | 23094 | -34.346 | <.0001 |
|  | WT TP1 - WT TP4 | -0.34797 | 0.00703 | 23094 | -49.504 | <.0001 |
|  | WT TP2 - WT TP3 | 0.06208 | 0.00703 | 23094 | 8.832 | <.0001 |
|  | WT TP2 - WT TP4 | -0.04447 | 0.00703 | 23094 | -6.327 | <.0001 |
|  | WT TP3 - WT TP4 | -0.10655 | 0.00703 | 23094 | -15.158 | <.0001 |
| **Within (Across Timepoints)**<br>**KO** | KO TP1 - KO TP2 | -0.19982 | 0.00703 | 23094 | -28.427 | <.0001 |
|  | KO TP1 - KO TP3 | -0.20552 | 0.00722 | 23094 | -28.462 | <.0001 |
|  | KO TP1 - KO TP4 | -0.15346 | 0.00703 | 23094 | -21.832 | <.0001 |
|  | KO TP2 - KO TP3 | -0.00570 | 0.00722 | 23094 | -0.790 | 0.4625 |
|  | KO TP2 - KO TP4 | 0.04636 | 0.00703 | 23094 | 6.595 | <.0001 |
|  | KO TP3 - KO TP4 | 0.05206 | 0.00722 | 23094 | 7.210 | <.0001 |

**Notes:** TP = timepoint (1 = 4 weeks, 2 = 8 weeks, 3 = 12 weeks, 4 = 15 weeks); KO = *Fmr1*<sup>-/-</sup> rats; WT = wild type rats; Positive *t*-value indicates higher functional connectivity in the first group; negative *t*-value indicates lower functional connectivity. Multiple comparisons were FDR corrected.

### Altered RSC connectivity in individuals with Fragile X Syndrome

To assess the translational relevance of our rat findings, we performed rsfMRI in a small cohort of young adults with FXS and age-matched typically developing controls (mean age 18.6 ± 0.5 years; n = 5 per group), focusing on the RSC as an a priori region of interest based on alterations observed in *Fmr1^-/y^* adult rats. Whole-brain DC analysis revealed trends toward reduced DC in the RSC and increased DC in the cerebellar vermis and insula in individuals with FXS compared to controls (Fig. 7A; p < 0.05, uncorrected), although these effects did not survive correction for multiple comparisons. Seed-based FC analysis using the RSC region revealed significantly reduced brain-wide RSC connectivity in individuals with FXS relative to controls (Fig.7B-D), a pattern consistent with the hypoconnectivity observed in adult *Fmr1^-/y^* rats.

**Fig. 7.**
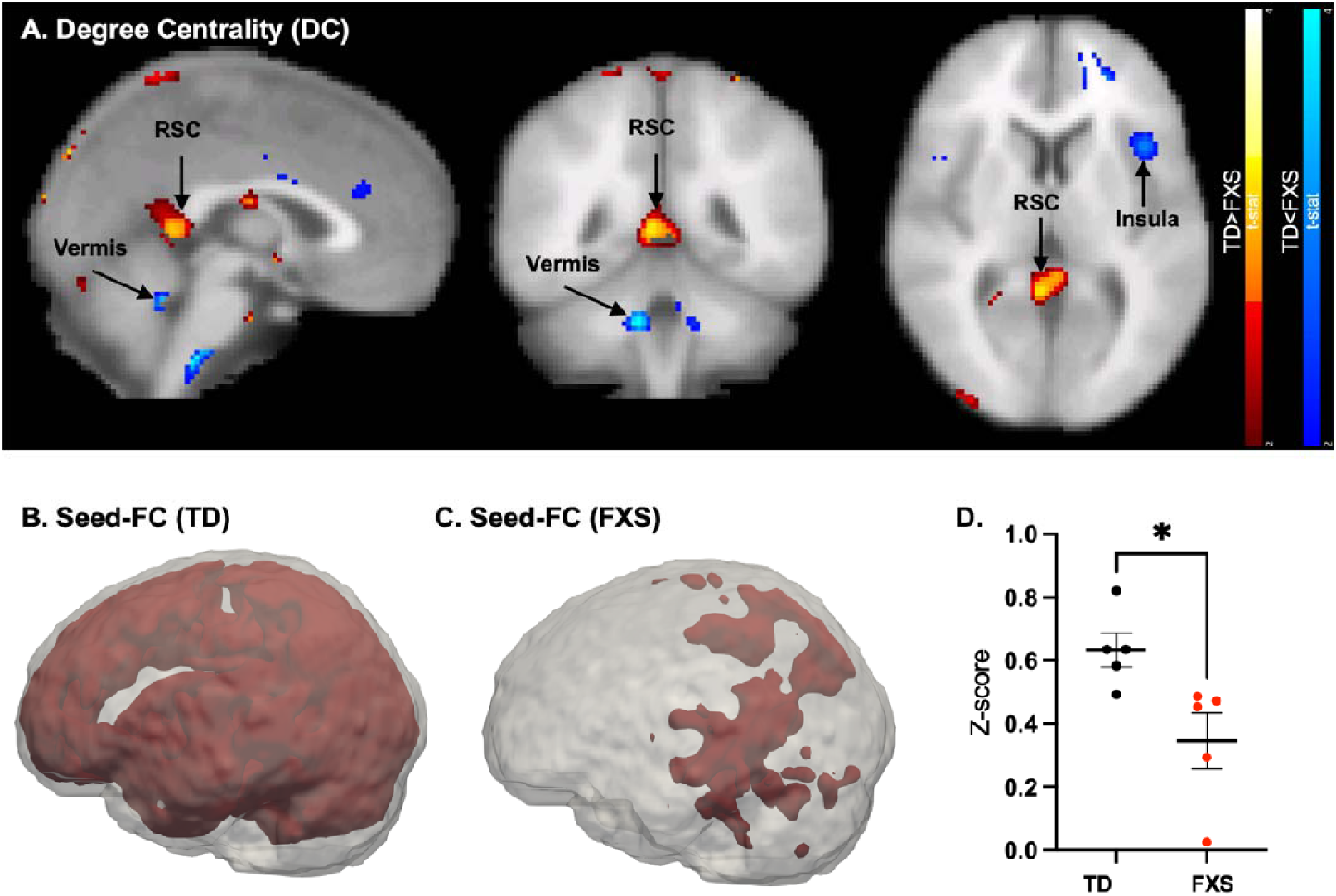
Reduced degree centrality and functional connectivity at RSC in individuals with FXS. (**A**) Degree centrality (DC) maps showing reduced DC in the RSC of individuals with FXS compared to typically developing (TD) controls. (**B, C**) Group-averaged RSC seed-based FC maps for TD controls (B) and individuals with FXS (C), showing widespread reduction in FC in the FXS group. Maps are displayed at r > 0.5. (**D**) Group comparison of RSC FC z-scores showing lower connectivity in individuals with FXS relative to TD controls (two-sample t-test, p < 0.05, T = 2.799). Data are shown as mean ± SEM. Group sizes: TD controls: n = 5; FXS: n = 5. Abbreviations: FXS, Fragile X syndrome; RSC, retrosplenial cortex; FC, resting functional connectivity; SEM, standard error mean.

## DISCUSSION

In this study, we used a longitudinal approach to determine how loss of FMRP affects the development of large-scale functional brain networks and whether these circuit-level abnormalities are sensitive to early-life pharmacological intervention. Loss of FMRP in rats altered the developmental trajectory of large-scale FC, with RSC hyperconnectivity in juveniles and widespread hypoconnectivity in adulthood. Early lovastatin treatment failed to normalize these network abnormalities despite prior evidence that the same intervention improves behavioural and cellular endpoints in this model (*12*). Instead, lovastatin shifted WT FC toward a *Fmr1^-/y^*-like profile, suggesting an unanticipated effect of treatment on large-scale network organization in WT animals. Together, these findings identify altered maturation of large-scale brain networks as a distinct feature of Fragile X-related pathophysiology and highlight the potential value of longitudinal circuit-level imaging for defining developmental trajectories and evaluating therapeutic effects.

A developmental framework is particularly important for interpreting circuit dysfunction in neurodevelopmental disorders, where altered trajectories may be more informative than abnormalities observed at single timepoints. Defining when network-level alterations emerge, and how they diverge from typical developmental trajectories, is therefore critical for understanding pathophysiological mechanisms and therapeutic opportunities. At 4 weeks of age, global resting-state network organization was broadly comparable between *Fmr1^-/y^*and WT rats, consistent with cross-species evidence that large-scale functional networks are established early in development (*27*). However, seed-based analyses revealed localized hyperconnectivity involving the RSC in *Fmr1^-/y^*rats, suggesting that early alterations were regionally specific rather globally distributed. Across development, FC increased in WT rats, whereas *Fmr1^-/y^* rats failed to follow this trajectory and instead exhibited a late reduction in connectivity by 15 weeks relative to earlier developmental stages. This divergence resulted in widespread genotype differences becoming apparent in adulthood. This developmental pattern suggests altered network maturation in *Fmr1^-/y^* rats, rather than a simple progressive loss of connectivity, with early regional alterations preceding broader disruptions in network organization. Similar age-dependent shifts in connectivity have been reported in FXS and ASD (*28*), supporting the concept that atypical developmental trajectories, rather than static network abnormalities, are a key feature of these conditions.

The developmental nature of these abnormalities is consistent with established roles for FMRP in synaptic refinement, experience-dependent plasticity, and critical-period regulation (*11, 29–32*). The 4-week timepoint coincides with a period of active circuit maturation in rodents, raising the possibility that early RSC hyperconnectivity reflects impaired activity-dependent pruning and/or stabilization of long-range projections. Similar circuit-dependent abnormalities have been reported in *Fmr1^-/y^*mice, albeit with regionally heterogeneous effects (*33–35*). Consistent with this framework, our human FXS cohort also suggested altered RSC connectivity, providing partial translational support for this regional phenotype.

The RSC is a particularly relevant node because of its role as a hub within the DMN (*36*) and its highly interconnected architecture linking hippocampal, cortical and sensory systems (*37*). This architecture may make the RSC sensitive to alterations in developmental circuit refinement, potentially contributing to DMN abnormalities reported in ASD and FXS (*38*). Consistent with this possibility, RSC-localized FMRP degradation in *Senp1*-deficient mice produces social and synaptic abnormalities that are rescued by local FMRP reintroduction (*39*). Although mechanistically distinct from global *Fmr1* deletion, this model underscores the sensitivity of the RSC to FMRP-dependent dysfunction. Human FXS studies likewise report DMN abnormalities, including altered connectivity in younger individuals and associations with behavioural severity and longitudinal progression (*8–10, 40*). These findings suggest that DMN dysfunction extends beyond the RSC, supporting the potential translational relevance of the rodent data.

The ERK signaling pathway has been implicated as a key mechanism disrupted in FXS, with loss of FMRP associated with dysregulation of Ras–ERK-dependent signaling and excessive protein synthesis (*41*). Consistent with this mechanism, lovastatin has been proposed to exert therapeutic effects, at least in part, through modulation of this pathway and has been shown across rodent models of FXS to rescue abnormalities in protein synthesis, synaptic plasticity, neuronal excitability, seizure susceptibility, and associative learning (*12, 14, 15*). In our previous work, the same treatment regimen used here prevented associative learning deficits in *Fmr1^-/y^*rats (*12*). These behavioural processes depend on coordinated activity across distributed networks, including prefrontal, hippocampal, and lateral entorhinal regions (*42–44*), suggesting that lovastatin can modify aspects of FMRP-dependent dysfunction relevant to circuit function. However, lovastatin did not rescue resting-state FC abnormalities in *Fmr1^-/y^* rats, which remained indistinguishable from untreated *Fmr1^-/y^* animals. These findings indicate that restoration of molecular, synaptic, and behavioural phenotypes reported following lovastatin treatment is not necessarily accompanied by restoration of intrinsic large-scale FC. Importantly, this does not imply that lovastatin lacks effects on circuit function, as resting-state connectivity captures only one dimension of network organization. Thus, our findings do not exclude the possibility that lovastatin restores local synaptic function, task-dependent recruitment of distributed networks, or other aspects of specific circuit processes without restoring the developmental trajectory of intrinsic large-scale connectivity.

Our findings may help explain the mixed outcomes observed in clinical studies of lovastatin in individuals with FXS (*16–18*), and also illustrate the challenges of predicting clinical efficacy from improvements observed at molecular, cellular, or behavioural levels in preclinical models. Our findings suggest a possible circuit-level explanation for this limited translation: improvements in molecular and behavioural phenotypes may not reflect widespread restoration of network function. These findings underscore the potential value of incorporating circuit-level measures as a physiologically distinct domain within therapeutic evaluation. Such an approach aligns with emerging multidomain frameworks, including the multidomain responder index (MDRI), which integrate clinically meaningful changes across independent domains within individuals to better capture treatment effects in complex and heterogeneous disorders (*45*).

Interestingly, lovastatin treatment instead appeared to shift FC in WT rats towards a more *Fmr1^-/y^*-like profile. Although not a primary aim of the study, this unexpected finding raises the possibility that the effects of lovastatin may differ depending on FMRP expression status. This may be particularly relevant for FXS, where X-chromosome- and incomplete *FMR1* inactivation results in mosaic FMRP expression, with FMRP-deficient and FMRP-expressing cells coexisting in the same individual. The effects of lovastatin in individuals with mosaic FMRP expression may therefore be complex and may not be predicted fully by studies of uniformly FMRP-deficient models. Further studies will be required to determine the basis and reproducibility of these effects. More broadly, these findings highlight that pathway modulation may have complex and context-dependent effects across levels of brain organization, which may contribute to variability in therapeutic outcomes.

Several limitations should be considered when interpreting these findings. The human cohort was modest in size and focused on a single developmental stage (adolescence/young adulthood). While they do highlight the importance of fMRI as a translational tool, replication across larger cohorts and across the lifespan will be important to determine the generalizability of these network-level alterations and to establish how they evolve with development in individuals with FXS. Preclinical imaging was performed under light anesthesia; although anesthesia can influence BOLD dynamics, resting-state networks remain detectable under the conditions used here (*46–48*). In addition, the MRI acquisition did not cover the entire rat posterior cerebellum, limiting our ability to detect connectivity alterations in this region. Finally, while the *Fmr1^-/y^*rat model provides a robust system for studying the consequences of complete FMRP loss, it does not capture the phenotypic variability in individuals with FXS. Together, these considerations motivate further investigation of circuit-level alterations across diverse FXS populations.

For clinical translation, future studies should determine whether RSC/DMN connectivity trajectories predict behavioural outcomes or treatment response longitudinally, and whether interventions targeting specific developmental windows or circuit mechanisms provide greater therapeutic benefit. Validation across independent cohorts, imaging platforms, and analytic pipelines will be essential to establish whether resting-state connectivity measures can serve as reliable biomarkers for treatment stratification or therapeutic efficacy in FXS. Future therapeutic studies incorporating circuit-level measures alongside molecular, cognitive, and behavioural endpoints may help determine when and how interventions modify disease-relevant biology beyond symptomatic improvement. Integrating resting-state connectivity measures with electrophysiological signatures, including EEG-based biomarkers of network excitability and synchrony (e.g. (*49*)), may further strengthen interpretation of circuit dysfunction and provide complementary measures of disease biology.

## MATERIALS AND METHODS

### Study design

A translational design was used to assess RSC-centered and global FC alterations in FXS. In rats, rsfMRI was performed longitudinally in male *Fmr1^-/y^*and WT littermates. In humans, rsfMRI was used to assess RSC-centered FC in individuals with FXS and age-matched typically developing controls.

### Rats

Male Long-Evans *Fmr1^-/y^* rats (LE-*Fmr1^em1Sidb^*) and WT littermates were bred in-house and maintained on a 12-hour light/dark cycle with food and water available ad libitum. Colony founders were generated by Horizon Discovery (formerly SAGE) using zinc-finger nuclease-mediated disruption of *Fmr1* with a targeted construct containing coding sequence for eGFP; resulting founders did not express FMRP or eGFP (*12*). Rats were housed in mixed-genotype cages (three to six per cage) and genotyped by PCR. All procedures were conducted in accordance with University of Edinburgh and UK Home Office regulations under the Animals (Scientific Procedures) Act 1986.

### Preclinical study design

> Rats were assigned to four groups: WT control diet, WT lovastatin diet, *Fmr1^-/y^* control diet, and *Fmr1^-/y^*lovastatin diet. Group sizes were n = 13, 14, 13, and 12, respectively. Mixed-genotype cages were randomly allocated to treatment conditions, and experimenters were blinded to genotype and treatment. Animals received standard chow ad libitum until P28, when they were switched to either control or lovastatin-enriched diet (100 mg/kg; Bio-Serv, USA). At P64, animals were returned to standard chow. Animals underwent resting-state fMRI at 4, 8, 12, and 15 weeks of age. Animal weights were monitored following the diet switch to ensure no adverse effects on development.

### Human Participants

Human participants were recruited through the Fragile X Society and Patrick Wild Centre research mailing lists. Ethical permission for the study was granted by Scotland A Research Ethics Committee. Where participants were unable to consent for themselves, consent was sought from their appropriate proxy (parent or guardian if under 16 years old; welfare guardian or nearest relative if over 16 years old, in accordance with the Adults With Incapacity (Scotland) Act 2000). Participants’ demographics are given in Table 3.

**Table 3.** Post-hoc contrasts of global rsFC across genotype and treatment groups at each timepoint.

| Timepoint | Contrast | Estimate | SE | t Ratio | p Value |
| --- | --- | --- | --- | --- | --- |
| **8 weeks** | KO CON – WT CON | -0.0575 | 0.0460 | -1.251 | 0.3254 |
|  | KO CON – KO LOVA | 0.00972 | 0.0470 | 0.207 | 0.8368 |
|  | KO CON – WT LOVA | -0.0695 | 0.0451 | -1.539 | 0.3174 |
|  | WT CON – KO LOVA | 0.0672 | 0.0470 | 1.432 | 0.3174 |
|  | WT CON – WT LOVA | -0.0120 | 0.0451 | -0.265 | 0.8368 |
|  | KO LOVA – WT LOVA | -0.080 | 0.0461 | -1.716 | 0.3174 |
| **15 weeks** | KO CON – WT CON | -0.1483 | 0.0460 | -3.227 | **0.0135** |
|  | KO CON – KO LOVA | -0.0230 | 0.0469 | -0.491 | 0.7509 |
|  | KO CON – WT LOVA | -0.0339 | 0.0451 | -0.752 | 0.6837 |
|  | WT CON – KO LOVA | 0.1253 | 0.0469 | 2.671 | **0.0291** |
|  | WT CON – WT LOVA | 0.1144 | 0.0451 | 2.535 | **0.0291** |
|  | KO LOVA – WT LOVA | -0.0109 | 0.0461 | -0.237 | 0.8140 |

**Notes:** KO = *Fmr1*<sup>-/-</sup> rats; WT = wild type rats; CON = control diet; LOVA = lovastatin diet. Positive *t*-value indicates higher functional connectivity in the first group; negative *t*-value indicates lower connectivity. Multiple comparisons were FDR corrected.

Given that individuals with FXS commonly experience anxiety in new environments, auditory hypersensitivity and hyperactivity, considerable preparation was undertaken prior to the scans to acclimatize the individuals to the scanning environment. This was done with the aid of two mock scanners (Fig. S3A). First, the individuals rehearsed on a crude replica of scanner (table, bore and head coil), taking as many visits as was necessary to be able to lie still in the mock scanner whilst listening to a recording of the scan sequence. Following this, participants were able to rehearse on a full-size replica of the real scanner prior to their research scan.

### Preclinical magnetic resonance imaging

Data acquisition was performed under light (1.5%) isoflurane anesthesia, maintained throughout the scanning session. Animal physiology was closely monitored throughout the session. Body temperature was monitored using a rectal probe and maintained at 37°C using a hot air ventilator. Breathing rate and oxygenation were monitored using a transducer pad and a pulse oximeter, respectively (Model 1030 monitoring and gating system, Small Animal Instrument Inc. Stony Brook, NY, USA).

Animals were positioned in a two-channel phased-array surface head coil (Rapid Biomedical, Rimpar, Germany) in a 7T horizontal bore Biospec ADVANCE neo preclinical imaging system, equipped with a 116 mm bore gradient insert, 660 mT/m gradient and 86 mm quadrature volume coil (Bruker BioSpin MRI GmbH, Ettlingen, Germany). A mouse head coil was used for juvenile rats (4 weeks of age) and a rat head coil was used for older animals (8 weeks and above). Resting state images were acquired using a one-shot gradient-echo Echo-Planar Imaging (GE-EPI) sequence (TR = 2000ms, TE = 12ms, in-plane resolution = 0.47 x 0.47mm, number of slices = 24, slice thickness = 0.85mm, slice gap = 0, volume = 300). Structural images were acquired using a T2-weighted RARE sequence (TR = 2310ms, TE = 36ms).

### Clinical magnetic resonance imaging

Data acquisition was performed on a 3T Siemens Magnetom Verio scanner. Functional resting state images were acquired using a single-shot EPI sequence (TR = 1560ms, TE = 26ms, in-plane resolution = 3.4375 x 3.4375mm, number of slices = 26, slice thickness = 5mm, volume = 293). Structural images were acquired using a T1-weighted magnetization prepared rapid acquisition gradient echo (MPRAGE) sequence (TR = 2300ms, TE = 2.98ms). Individuals were asked to remain awake with their eyes open and to focus on a fixation cross seen through goggles in the scanner.

### Resting state fMRI data analysis

#### Preprocessing

rsfMRI datasets were first pre-processed using Statistical Parametric Mapping (SPM12; Wellcome Department of Imaging Neuroscience, London, UK). Preclinical structural and functional images were scaled by a factor of 10 in the x, y and z direction to account for size difference between the rat and human brain. The first 10 volumes of each 4D functional image were discarded to allow for T1 equilibration effects. Skull-stripping was performed using in-house MATLAB (The Mathworks Inc, version 2020a) scripts for rodent images and using FSL’s Brain Extraction Tool (BET) for clinical images. The datasets were registered to a high-resolution anatomical template (SIGMA rat brain template for rat data (*50*), MNI template for human data) using SPM12. An 8mm FWHM spatial smoothing was then applied, and bandpass filtering [0.01 0.1 Hz] was performed to preserve low-frequency fluctuations in the resting state BOLD signal. One 8-week-old and one 12-week-old rat were excluded following preprocessing due to excessive head motion (> 1 voxel size); both rats belonged to the *Fmr1^-/y^* group. Representative ICA-derived resting-state networks from the rat and human datasets, together with representative human motion estimates, are shown in Figs. S2 and S3B–D, respectively.

#### Degree Centrality (DC)

DC was computed using DPABI (*51*). For each voxel, Pearson correlation coefficients were calculated between its time series and that of every other voxel in the brain. A correlation threshold of r > 0.25 was applied to suppress spurious connections (*20*). The weighted degree of each voxel was calculated as the sum of suprathreshold correlations. These voxel-wise DC values were then normalized by dividing by the global mean DC value, producing standardised DC maps. Finally, these maps were spatially smoothed using a 4 mm full width at half maximum (FWHM) Gaussian kernel.

Seed-based FC analysis: In seed-based FC analysis, the BOLD time series was extracted from the average of a defined seed region of interest, and Pearson correlation coefficients were computed between this signal and all other voxels in the brain. A correlation threshold of r > 0.5 was applied to define meaningful functional connections, in line with prior work identifying this cutoff as indicative of moderate and behaviourally relevant connectivity in rsfMRI (*52, 53*). The resulting voxel-wise FC maps were then Fisher’s Z-transformed for group-level comparisons.

Seed-based within-network FC analysis: The DMN was identified using ICA of resting-state fMRI data (Fig. S2). The resulting ICA-derived DMN spatial map was thresholded to generate a binary DMN mask. To quantify RSC integration within the DMN, a seed-to-voxel FC analysis was performed using the RSC seed defined from the prior DC analysis. Pearson correlation coefficients (r) were calculated between the mean BOLD time series of the RSC seed and the BOLD time series of each individual voxel within the DMN mask. The resulting voxel-wise correlation values were converted to z-scores using Fisher’s z transformation and averaged across all mask voxels, yielding a single measure of RSC-DMN connectivity strength for each participant.

#### ROI-based FC analysis

BOLD time series were extracted and averaged from 25 anatomically defined brain regions using a custom ROI atlas (Fig. 2A) generated by combining regions from the Fisher 344 and SIGMA rat brain atlases (*21, 50*). Pairwise Pearson correlation coefficients were computed between all ROI time series, resulting in a symmetric FC matrix for each subject. These correlation matrices were Fisher-Z transformed for statistical comparison.

### Statistical analysis

Group-level differences were assessed using two-sample t-tests and mixed-effects models. ROI-to-ROI pairwise comparisons were corrected for multiple comparisons using network-based statistic (NBS; 1,000 permutations)(*54*). Voxel-wise comparisons were corrected using threshold-free cluster enhancement (TFCE; p < 0.05)(*55*).

For pairwise comparison of longitudinal rat data, dimensionality was reduced by vectorizing each subject’s FC matrix (25 × 25 ROIs) into 300 unique values, which were then analysed with linear mixed-effects (LME) models in R (*56*). The model was corrected for multiple connections of interest and was hierarchically structured with random intercepts and network connections nested within animals, thus including all network connections into a single model. Consequently, the output only informs on statistical significance for the whole model, and not for any individual connections.

## Acknowledgments

We thank R. Lennen, M. Jansen, and M. Walls for assistance in fMRI data collection and colleagues in the Patrick Wild Centre and Simons Initiative for the Developing Brain for constructive discussions during the course of this study. BVS for veterinary advice and animal husbandry/maintenance. Animal technicians: W. Mungall, R. Shiels, A. Onishi,

## Author contributions

Conceptualization: PCK SMT AS

Methodology: JY, JS, AM, MS

Investigation: JS, AM

Visualization: JY, JS, AM

Funding acquisition: PCK, AS, AM, SMT

Project administration: JS, SMT

Supervision: SMT, PCK, AS, RD

Writing – original draft: JY, JS, SMT, AM

Writing – review & editing: PCK, JY, SMT, others

## Data and materials availability

Data that support the findings of this study are available from the authors upon reasonable request. Scripts used in this study will be available on Github.

## List of Supplementary

### Materials Materials and Methods

#### Independent component analysis (ICA)

Following preprocessing, resting-state networks (RSNs) were identified using spatial independent component analysis (ICA) as implemented in FSL MELODIC (*57*). Group-level ICA was performed independently for the rat and human datasets using a temporal concatenation approach. For each dataset, a low-dimensional decomposition was applied to extract 15 independent components (ICs). This dimensional constraint was chosen to optimize the identification of major large-scale networks while avoiding excessive component splitting. The resulting ICs were classified as signal or noise based on established spatiotemporal criteria, including core structural overlap, low-frequency spectral dominance, and the absence of edge-focused motion artifacts (*58*).

To ensure unbiased network identification across species, signal components were matched to canonical RSN templates using quantitative spatial cross-correlation:

Human Network Identification: Signal components were spatially correlated against standard canonical resting-state maps (*59*).

Rat Network Identification: Signal components were spatially correlated against rat resting-state network template (*60*).

Representative IC maps for rats and humans are presented in Fig. S2 and Fig. S3D, respectively.

#### Principle component analysis (PCA)

To reduce the dimensionality of resting-state FC matrices prior to longitudinal analysis, PCA was performed on the vectorised upper triangle of each subject’s FC matrix. The first three principal components (PCs), capturing the greatest sources of variance across subjects, were retained. Subject-specific loadings/scores on each PC were extracted and entered as dependent variables in linear mixed-effects models, with timepoint as a fixed effect and subject as a random effect, to test for longitudinal changes in whole-matrix connectivity patterns whilst accounting for repeated-measures structure.

---

**Fig. S1.**
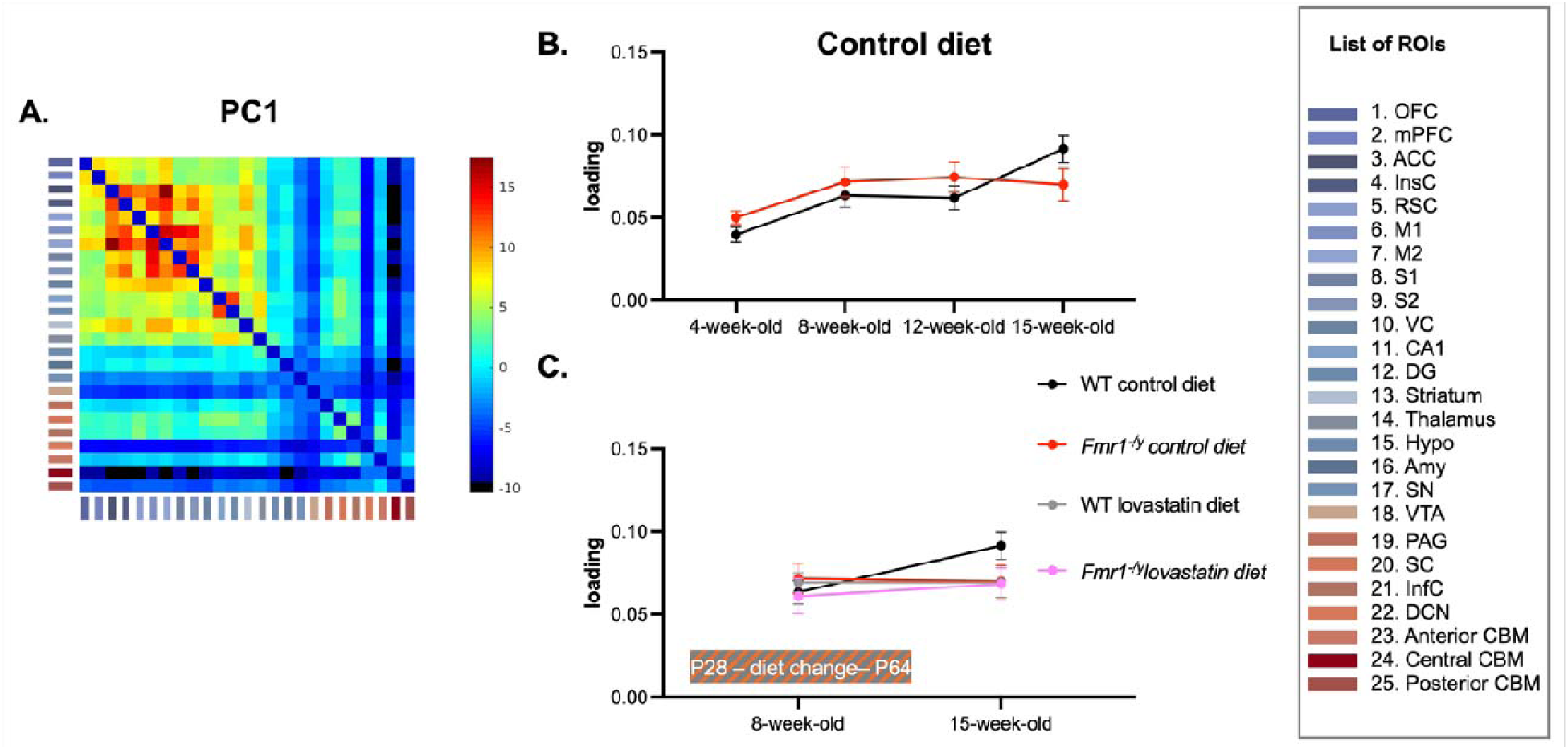
Principal component analysis (PCA) of FC changes across development and following lovastatin treatment. (A) The first component (PC1), accounting the greatest proportion of variance across global connectivity matrices. (B) PC1 loadings across development for WT and *Fmr1^-/y^* rats receiving control diet. (C) PC1 loadings during (8 weeks) and following (15 weeks) lovastatin treatment in for WT and *Fmr1^-/y^*rats. Data are shown as mean ± SEM. Group sizes: WT control diet, n = 13; *Fmr1^-/y^*control diet, n = 13; WT lovastatin diet, n = 14; *Fmr1^-/y^* lovastatin diet, n = 11 (8 weeks) and n = 12 (15 weeks). Mixed-effects analysis. B: main effect of genotype, F_(1,24)_ = 0.12, p = 0.73; main effect of age, F_(3,71)_ = 9.339, p < 0.0001; genotype × age interaction, F_(3,71)_ = 2.654, p = 0.0551. C: main effect of genotype, F_(1,48)_ = 0.70, p = 0.41; main effect of age, F_(1,47)_ = 2.57, p = 0.12; main effect of treatment, F_(1,48)_ = 1.13, p = 0.29; genotype × treatment interaction, F_(1,48)_ = 0.02, p = 0.89; genotype × age interaction, F_(1,47)_ = 1.09, p = 0.30; treatment × age interaction, F_(1,47)_ = 0.77, p = 0.39; age × genotype × treatment interaction, F_1,47)_ = 3.09, p = 0.09. Abbreviations: SEM, standard error mean; WT wild-type.

**Fig. S2.**
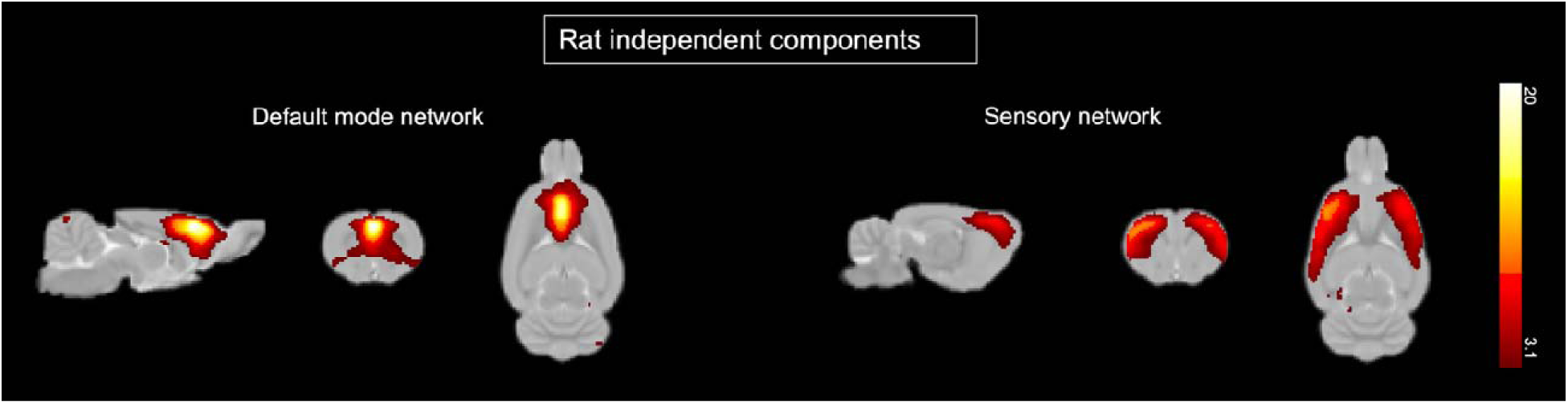
Representative independent components identified by independent component analysis (ICA) in 15-week-old rats. Left: default mode network; Right: sensorimotor network.

**Fig. S3.**
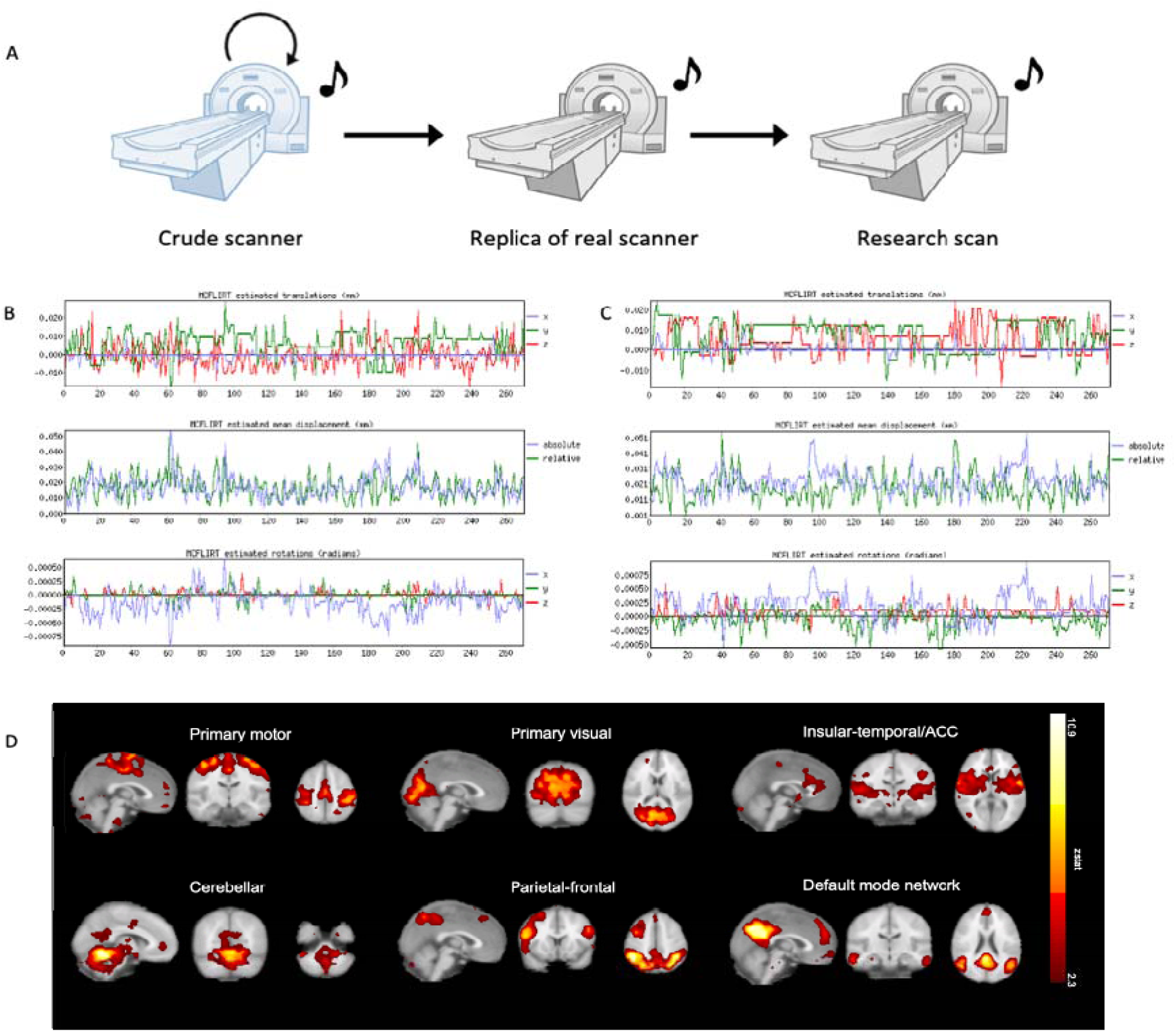
Resting state fMRI scanning in individuals with Fragile X Syndrome. (A) Schematic of acclimatization protocol for FX individuals. To habituate themselves to the scanning procedure, participants practice laying still on a crude replica of the scanner as many time as it took them to be comfortable. They next had a habituation session on an accurate replica of the scanner and finally had the research scan. (B) Representative motion traces of a typically developing individual. From top-to-bottom, traces of the estimated translation in mm in the x, y, and z directions, of the absolute and relative estimated mean displacement in mm and of the estimated rotation in radians in the x, y and z directions. (C) Representative motion traces of a FX individual. From top-to-bottom, traces of the estimated translation in mm in the x, y, and z directions, of the absolute and relative estimated mean displacement in mm and of the estimated rotation in radians in the x, y and z directions. (D) Resting State Networks (overlaid on a high-resolution anatomical template shown in the sagittal, coronal and axial planes) in individuals with severe FXS, identified by ICA. From left to right, upper to lower: Primary Motor Network, Primary Visual Network, Insular-Temporal / ACC Network, Cerebellar Network, Parietal-Frontal Network, Default Mode Network. Colour bars represent Z-score, ranging from 4-43.

